# Characterization of Tigurilysin, a Novel Human CD59-Specific Cholesterol-Dependent Cytolysin, Reveals a Role for Host Specificity in Augmenting Toxin Activity

**DOI:** 10.1101/2023.06.21.545930

**Authors:** Ifrah Shahi, Sophia A. Dongas, Juliana K. Ilmain, Victor J. Torres, Adam J. Ratner

## Abstract

Cholesterol dependent cytolysins (CDCs) are a large family of pore forming toxins, produced by numerous gram-positive pathogens. CDCs depend on host membrane cholesterol for pore formation; some CDCs also require surface associated human CD59 (hCD59) for binding, conferring specificity for human cells. We purified a recombinant version of a putative CDC encoded in the genome of *Streptococcus oralis subsp. tigurinus*, tigurilysin (TGY), and used CRISPR/Cas9 to construct hCD59 knockout (KO) HeLa and JEG-3 cell lines. Cell viability assays with TGY on WT and hCD59 KO cells showed that TGY is a hCD59-dependent CDC. Two variants of TGY exist among *S. oralis subsp. tigurinus* genomes, only one of which is functional. We discovered that a single amino acid change between these two TGY variants determines its activity. Flow cytometry and oligomerization western blots revealed that the single amino acid difference between the two TGY isoforms disrupts host cell binding and oligomerization. Furthermore, experiments with hCD59 KO cells and cholesterol depleted cells demonstrated that TGY is fully dependent on both hCD59 and cholesterol for activity, unlike other known hCD59-dependent CDCs. Using full-length CDCs and toxin constructs differing only in the binding domain, we determined that having hCD59-dependence leads to increased lysis efficiency, conferring a potential advantage to organisms producing hCD59-dependent CDCs.

**IMPORTANCE:** Cholesterol dependent cytolysins (CDCs) are produced by a variety of disease-causing bacteria, and may play a significant role in pathogenesis. Understanding CDC mechanisms of action provides useful information for developing anti-virulence strategies against bacteria that utilize CDCs and other pore-forming toxins in pathogenesis. This study describes for the first time a novel human-specific CDC with an atypical pore forming mechanism compared to known CDCs. In addition, this study demonstrates that human-specificity potentially confers increased lytic efficiency to CDCs. These data provide a possible explanation for the selective advantage of developing hCD59-dependency in CDCs and the consequent host restriction.

## INTRODUCTION

Pore forming toxins (PFTs) are produced by many pathogenic bacteria and form a diverse class of bacterial protein toxins (1). PFTs likely contribute to bacterial pathogenesis in several ways, including enhancement of colonization (2–4), release of host cell nutrients (5), hijacking of host cell pathways (6, 7), and intracellular delivery of other virulence factors (8, 9). PFTs function by perforating cell surface membranes, often leading to osmotic lysis, or triggering other downstream effects, such as pro-inflammatory cell signaling cascades, membrane repair, or programmed cell death (10).

Cholesterol dependent cytolysins (CDCs) are a family of PFTs produced by many gram-positive bacteria (10, 11), as well as some gram-negative bacteria (12). CDCs are characterized by large pores and by a dependency on host membrane cholesterol (11, 13, 14). For most CDCs, the presence of cholesterol in eukaryotic cell plasma membranes is sufficient for targeting and lysing a cell (11). A subset of CDCs additionally requires human CD59 (hCD59), a cell surface molecule, for binding to host membranes.

CDC protein structures include four domains (15, 16). During pore formation, Domain 4 binds to the target cell membrane, anchoring the CDC to the cell surface. For most CDCs, this entails binding to host cell membrane cholesterol via a threonine-leucine pair conserved within the Cholesterol Recognition Motif (CRM), followed by insertion of several other structural loops including a conserved 11-amino acid sequence (undecapeptide [UDP]), providing anchorage and stability. For hCD59-dependent CDCs, the toxin first binds to hCD59 via a conserved sequence in Domain 4, before further being stabilized by cholesterol binding (16–18). Once a CDC is anchored to the host cell surface, >30 units oligomerize to form a beta-barrel pore (19–21). To date, four hCD59-dependent CDCs have been characterized: intermedilysin (ILY) produced by *Streptococcus intermedius*, lectinolysin (LLY) produced by *Streptococcus mitis*, vaginolysin (VLY) produced by *Gardnerella vaginalis*, and most recently, discoidinolysin (DLY) produced by *Streptococcus mitis* (17, 22–24). Since there are few established hCD59-dependent CDCs, it is difficult to form a comprehensive overview of the evolution of hCD59-dependency and how that contributes to virulence for the bacteria producing such CDCs.

*Streptococcus oralis subsp. tigurinus* is a commensal bacterium found in the human oral microbiome that has been implicated in invasive infections such as infective endocarditis, spondylodiscitis, and meningitis (25–28). It is an α-hemolytic, non-motile, and non-spore-forming bacterium. The *Streptococcus oralis subsp. tigurinus* genome includes an open reading frame with sequence similarity to other hCD59-dependent CDC genomes (17, 18). Using a recombinant protein, we characterized this novel hCD59-dependent CDC, tigurilysin (TGY). Of the two TGY alleles present in the NCBI database, one encodes a protein that is inactive at physiological concentrations, and we identified a single amino acid change in Domain 4 responsible for its activity. Furthermore, we delved into the question of hCD59-dependency and showed that while targeting cells with hCD59 restricts such CDCs to only one species, it also increases their lytic efficiency on epithelial cells by several orders of magnitude.

## RESULTS

### There are two variants of the *Streptococcus oralis subsp. tigurinus* CDC gene

Whole genome sequencing of three *Streptococcus oralis subsp. tigurinus* strains has revealed that some, but not all, strains have an open reading frame encoding a protein with similarity to previously characterized CDCs (29). We compared the amino acid sequences of *Streptococcus oralis subsp. tigurinus* CDC (tigurilysin [TGY]) from two strains: AZ_3a and AZ_14. Five amino acid differences were found between the mature predicted amino acid sequences of the two variants (**Fig 1A**). We also compared TGY with amino acid sequences from other CDCs. Previous studies with ILY have revealed that the toxin binds to hCD59 via residues separate from those of the cholesterol-recognizing motifs (17, 30). Comparison among CDC amino acid sequences has shown hCD59-binding residues to be conserved among hCD59-binding CDCs, and they are also found in the TGY amino acid sequence. Furthermore, the UDP of hCD59-binding CDCs harbors a proline in the 9^th^ position instead of a tryptophan (18); this proline is also seen in the TGY amino acid sequence. Based on these observations, we predicted that TGY was likely a hCD59-dependent toxin (**Fig 1B**).

**Figure 1:**
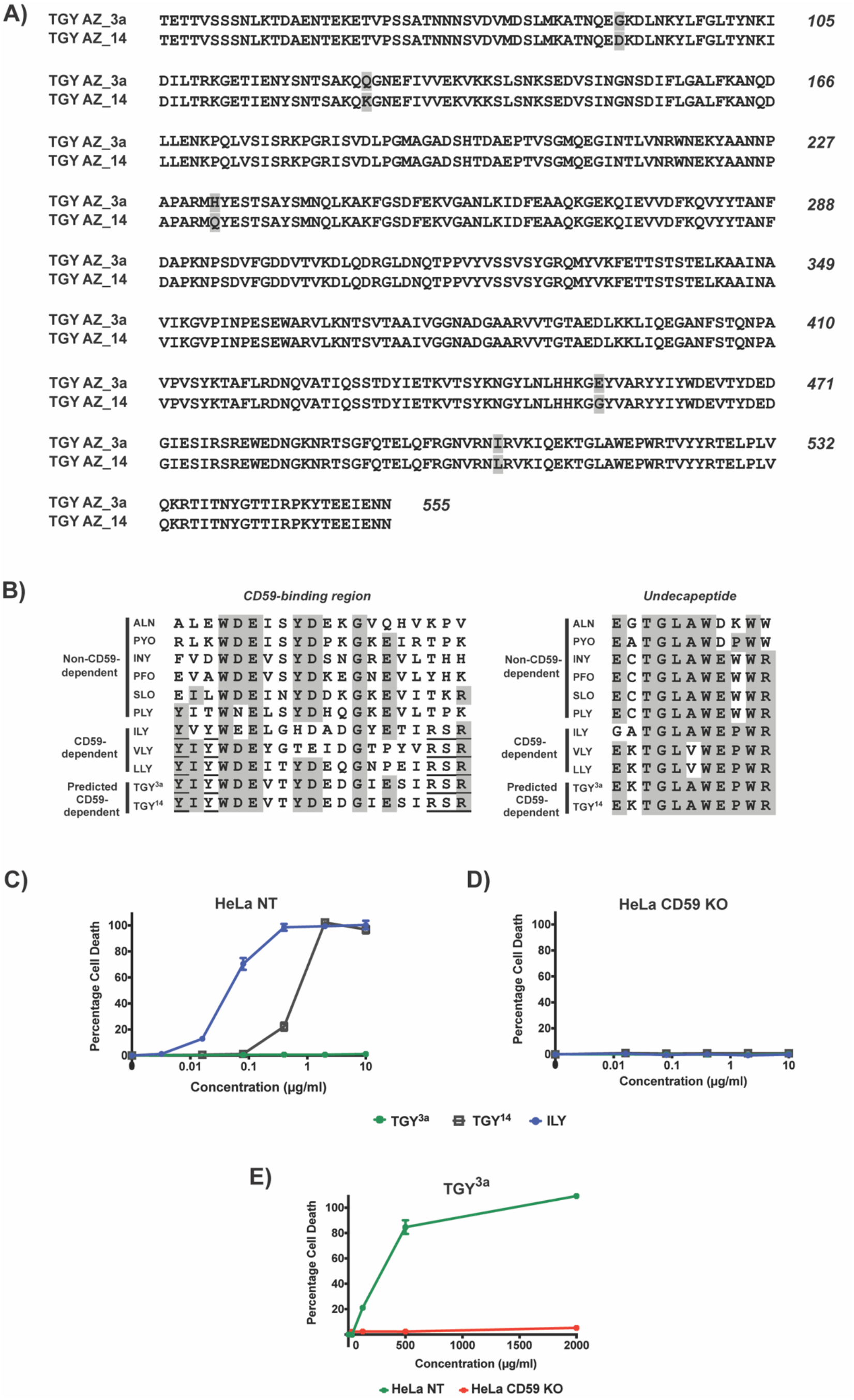
Two variants of novel CDC tigurilysin (TGY) exist, with differing lysis activity. (**A**) The predicted amino acid sequences of the two mature TGY variants. The signal sequence comprising the first 44 amino acids, as predicted by SignalP 5.0 (<u>SignalP 5.0 - DTU Health Tech -</u> <u>Bioinformatic Services</u>), was eliminated from the sequences shown here. There are 5 single amino acid differences (shaded) between TGY^3a^ and TGY^14^. The last two (amino acid 454 and amino acid 544) are found in domain 4. (**B**) Left panel: the conserved hCD59-binding sequence (Y-X-Y-X14-R-S-R; underlined) in Domain 4 of various hCD59-dependent CDCs, as compared to the same region of non-hCD59-dependent CDCs. Right panel: The conserved 11-amino acid sequence (undecapeptide), necessary for CDC interaction with cholesterol and subsequent pore formation, as found in various CDCs. The proline at the 9^th^ position is conserved among hCD59-dependent CDCs. The shaded regions in both panels show the consensus sequence as determined by MacVector. (**C-D**) Percentage cell death in HeLa NT control cells and HeLa hCD59 KO cells when exposed to increasing concentrations of TGY^3a^ and TGY^14^, with ILY for comparison as a known hCD59-dependent CDC. Cells were incubated with toxins for 1.5 hours and cell death measured by an LDH-release cytotoxicity assay. Each point is the mean of 3 replicates, and error bars represent ±SD. (**E**) Percentage cell death in HeLa NT control cells and HeLa hCD59 KO cells when exposed to supraphysical concentrations of TGY^3a^. Cells were incubated with toxin for 1.5 hours and cell death measured by an LDH-release cytotoxicity assay. Each point is the mean of 3 replicates, and error bars represent ±SD.

Recombinant His-tagged versions of TGY based on the sequences from both strains were created, expressed in *Escherichia coli*, and purified. In order to assess the role of hCD59 in host cell susceptibility to TGY, we utilized HeLa cells, a human cell line that expresses hCD59 (**Fig S1**). We used cells that were transduced with CRISPR-Cas9 single guide RNA targeting the hCD59 gene (HeLa hCD59 Knockout [KO]) and HeLa control cells that have been transduced with a non-targeting sgRNA (HeLa NT). We have previously used these cell lines to successfully examine ILY activity and hCD59-dependency (31). The two recombinant TGY variants were tested for lysis on the HeLa NT cells and HeLa hCD59 KO cells, and IC_50_ values (i.e. concentration required for 50% lysis of HeLa cells) were calculated. AZ_14 TGY (TGY^14^) lysed HeLa cells in a hCD59-dependent manner (IC_50_: 460 ng/ml), albeit at concentrations higher than those needed for lysis by ILY (IC_50_: 30 ng/ml). In contrast, the AZ_3a TGY (TGY^3a^) did not induce lysis at the concentrations tested, appearing to be inactive (**Fig 1C and 1D**). When TGY^3a^ was further tested on HeLa cells at supraphysiological concentrations, it was found to cause lysis in a hCD59-dependent manner, indicating that the toxin is functional but inefficient, requiring high dosages for functionality (IC_50_: 261 μg/ml) (**Fig 1E**). To confirm that this hCD59-dependent lysis is not specific to HeLa cells only, both TGY variants were also tested for lysis on JEG-3 cells (a human choriocarcinoma cell line) with and without hCD59 expression, with results similar to those on HeLa cells: TGY^14^ lysed the cells in a hCD59-dependent manner, while TGY^3a^ remained inactive at the concentrations tested (**Fig S2**).

### TGY residue at position 454 is important for lytic efficiency

To understand the cause of TGY^3a^ inefficiency, we turned our attention to the amino acid sequence differences between the two TGY variants. Of the five single amino acid differences, two are in Domain 4: residue 454 and residue 505 (**Fig 1A**). Residue 505 incorporates an isoleucine in TGY^3a^ and a leucine in TGY^14^. Based on amino acid sequences of other CDCs, this residue position seems to be permissible for both amino acids – some CDCs have an isoleucine while others have a leucine, without any correlation to the species specificity of the CDC. Conversely, residue 454 is part of the short hydrophobic loop known as Loop 2 (L2), one of the three conserved CDC loops that insert into the host membrane in a cholesterol-dependent manner (32, 33). In most CDCs, the corresponding amino acid in L2 is an alanine; however, non-functional TGY^3a^ harbors a glutamic acid (E) at this position, while the functional TGY^14^ has a glycine (G). Additionally, both TGY variants have an isoleucine (I) instead of a leucine (L) at position 544, which is part of the conserved CRM threonine-leucine pair.

Using site-directed mutagenesis, we made three amino acid substitutions in TGY^3a^, resulting in TGY^3a_E454G^, TGY^3a_E454A^ and TGY^3a_I544L^. These variants were tested on HeLa NT cells for lysis. The results demonstrated that the single amino acid change at position 454 restored the toxin’s lytic activity (**Fig 2A**), while the amino acid change in the CRM had no effect. TGY^3a_E454G^ and TGY^3a_E454A^ were further tested for lysis on HeLa hCD59 KO cells, confirming their cytotoxicity to still be hCD59-dependent (**Fig 2B**).

**Figure 2:**
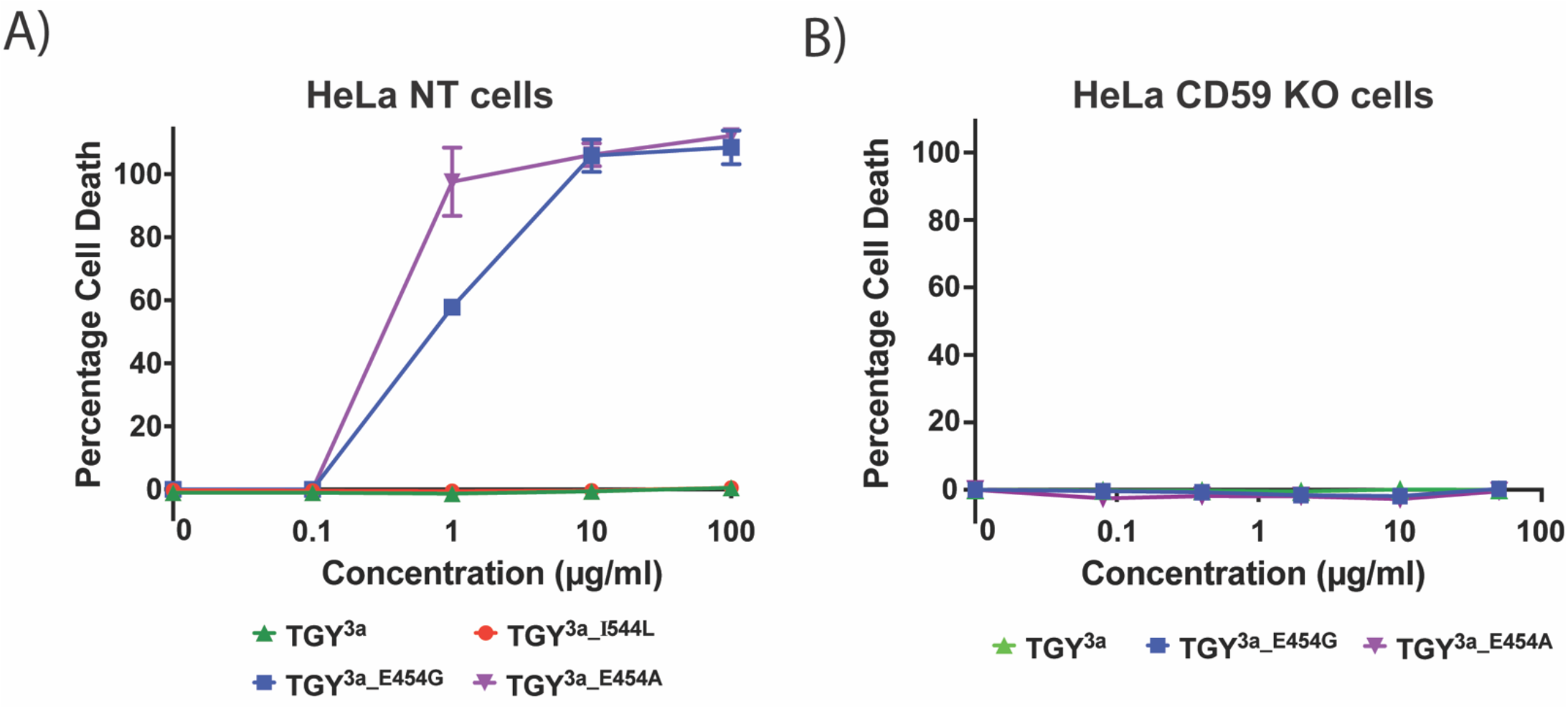
A glutamic acid-to-glycine change at position 454 affects TGY activity. (**A**) Percentage cell death in HeLa NT control cells when exposed to increasing concentrations of TGY^3a^ and its mutagenized counterparts. Only the amino acid change at position 454 (glutamic acid to glycine or alanine) restored toxin functionality. Cells were incubated with toxins for 1.5 hours and cell death measured by an LDH-release cytotoxicity assay. Each point is the mean of 3 replicates, and error bars represent ±SD. (**B**) Percentage cell death in HeLa hCD59 KO cells when exposed to increasing concentrations of TGY^3a^, TGY^3a_E454G^ and TGY^3a_E454A^, to determine if the toxins are hCD59-dependent. Cells were incubated with the toxins for 1.5 hours and cell death measured by an LDH-release cytotoxicity assay. Each point is the mean of 3 replicates, and error bars represent ±SD.

### TGY is fully dependent on both cholesterol and hCD59

Of the known hCD59-dependent CDCs, only ILY is completely non-functional in the absence of hCD59 – it requires hCD59 to initiate binding to a host cell, though interaction with cholesterol is still necessary for the subsequent steps of pore formation (34, 35). In contrast, VLY can bind host cell membrane cholesterol in the absence of hCD59 (though less efficiently than binding hCD59) (36), and LLY is similar (37). To assess whether TGY requires cholesterol in a manner similar to ILY or VLY/LLY, HeLa cells were depleted of cholesterol using methyl-ß-cyclodextrin (MßCD) and tested with TGY variants for lysis (**Fig 3A and 3B**). For comparison, similar assays were conducted with ILY (fully hCD59-dependent), VLY (partially hCD59-dependent), and pneumolysin (PLY; a non-specific CDC) (**Fig 3C**, **3D, and 3E**).

**Figure 3:**
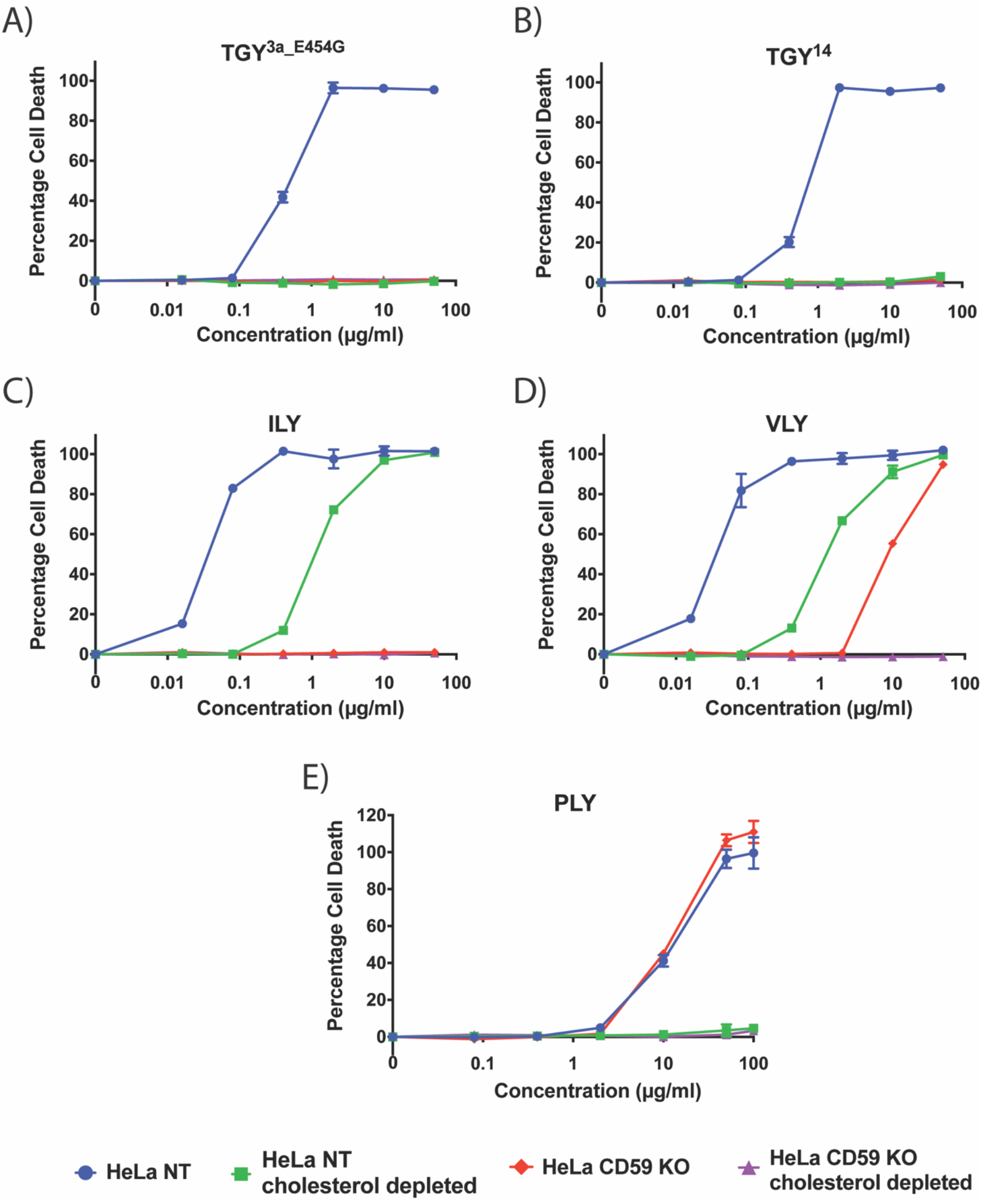
Functional TGY is fully cholesterol dependent. (**A-E**) Percentage cell death in HeLa NT control cells, HeLa hCD59 KO cells, HeLa cholesterol depleted cells, and HeLa hCD59 KO + cholesterol depleted cells. Cells were exposed to a range of concentrations of each toxin for 1.5 hours, and cell death measured by an LDH-release cytotoxicity assay. Each point is the mean of 3 replicates, and error bars represent ±SD.

Our results show that ILY does not lyse HeLa hCD59 KO cells but remains active on cholesterol depleted HeLa cells at higher toxin concentrations. VLY, as a partially hCD59-dependent CDC, lyses both HeLa hCD59 KO cells and cholesterol depleted HeLa cells at higher toxin concentrations. PLY, as a non-specific CDC, lyses HeLa hCD59 KO cells at similar levels as HeLa NT cells, but is inactive on cholesterol depleted HeLa cells. In contrast, TGY only lyses HeLa NT cells and does not kill cells either lacking hCD59 or depleted of cholesterol. To ensure that cholesterol depletion does not perturb hCD59 expression (and thereby indirectly affect lysis results of hCD59-dependent toxins), hCD59 cell surface expression was analyzed using flow cytometry (**Fig S1**). Results demonstrate that cholesterol depleted cells express similar levels of hCD59 as HeLa NT cells. Taken together with the lysis data, our results suggest that TGY has a distinct CDC lysis mechanism with an absolute requirement for both cholesterol and hCD59.

### TGY binding to host cells is affected by both cholesterol and hCD59 presence

We set out to investigate sequential stages of CDC pore formation for TGY, starting with host cell binding. HeLa cells were incubated with His-tagged TGY variants for flow cytometry analysis (**Fig 4A**). The same experiment was also carried out with ILY, VLY and PLY (**Fig 4B**). The percentage of PE-positive cells (indicating percentage of cells with detectable toxin bound to their surface) and the mean fluorescence intensity (MFI – indicating the density of toxin bound) for all conditions are listed in **Table 1**.

**Figure 4:**
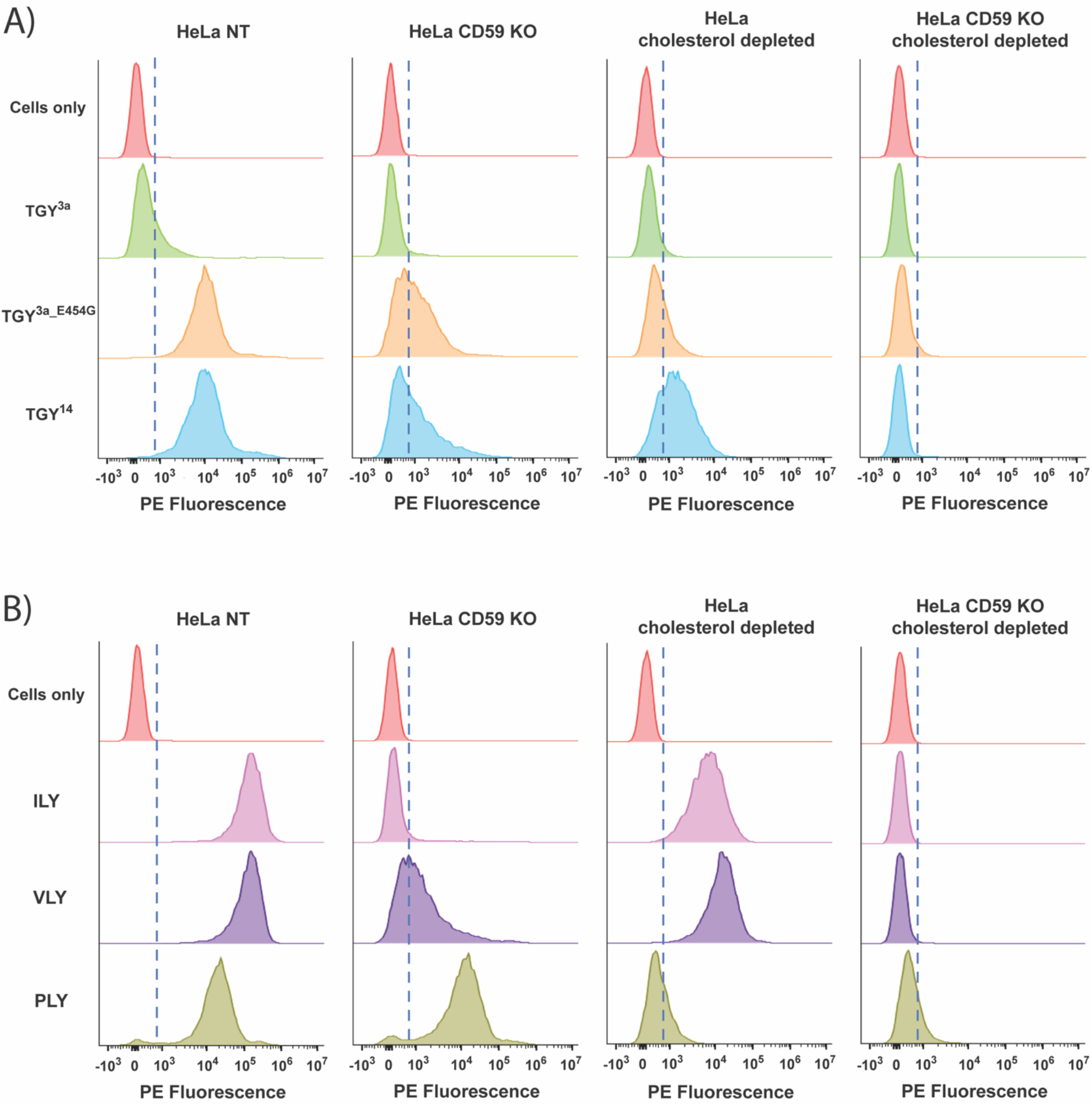
TGY3a is defective in host cell binding, while functional TGY is dependent on both cholesterol and hCD59 for host cell binding. (**A-B**) Flow cytometry analysis of His-tag expression on the cell surface after HeLa cells (NT control, hCD59 KO, cholesterol depleted, and hCD59 KO + cholesterol depleted) were incubated with His-tagged toxins, before addition of a PE anti-His-tag antibody. Histogram x-axis shows fluorescence intensity of the PE-conjugated antibody for each cell population. Histogram y-axis shows cell count normalized to mode. The dotted line demarcates the PE-negative cells and the PE-positive cells, based on the fluorescence intensity of control cells not incubated with any toxin.

**Table 1:** Quantification of Flow Cytometry Data of CDC Binding on HeLa cells. The percentage of PE-positive live cells were calculated by FlowJo based on gating of PE-negative cells as determined by cells-only controls. The MFI for each sample was calculated by FlowJo for cells gated as live cells → single cells.

|  | % PE-positive cells | MFI values |
| --- | --- | --- |
| HeLa NT cells only | 1 | 112 |
| HeLa hCD59 KO cells only | 1 | 578 |
| HeLa NT cholesterol depleted cells only | 0 | 136 |
| HeLa hCD59 KO cholesterol depleted cells only | 1 | 138 |
| HeLa NT + ILY | 100 | 170246 |
| HeLa hCD59 KO + ILY | 11 | 6767 |
| HeLa NT cholesterol depleted + ILY | 99 | 9910 |
| HeLa hCD59 KO cholesterol depleted + ILY | 1 | 151 |
| HeLa NT + VLY | 100 | 150828 |
| HeLa hCD59 KO + VLY | 65 | 8167 |
| HeLa NT cholesterol depleted + VLY | 100 | 21252 |
| HeLa hCD59 KO cholesterol depleted + VLY | 2 | 181 |
| HeLa NT + PLY | 96 | 30816 |
| HeLa hCD59 KO + PLY | 95 | 23677 |
| HeLa NT cholesterol depleted + PLY | 32 | 697 |
| HeLa hCD59 KO cholesterol depleted + PLY | 24 | 625 |
| HeLa NT + TGY<sup>3a</sup> | 28 | 9735 |
| HeLa hCD59 KO + TGY<sup>3a</sup> | 6 | 335 |
| HeLa NT cholesterol depleted + TGY<sup>3a</sup> | 7 | 4261 |
| HeLa hCD59 KO cholesterol depleted + TGY<sup>3a</sup> | 1 | 598 |
| HeLa NT + TGY<sup>3a</sup><sub>E454G</sub> | 99 | 28460 |
| HeLa hCD59 KO + TGY<sup>3a</sup><sub>E454G</sub> | 60 | 3813 |
| HeLa NT cholesterol depleted + TGY<sup>3a</sup><sub>E454G</sub> | 34 | 3847 |
| HeLa hCD59 KO cholesterol depleted + TGY<sup>3a</sup><sub>E454G</sub> | 8 | 281 |
| HeLa NT + TGY<sup>14</sup> | 99 | 31109 |
| HeLa hCD59 KO + TGY<sup>14</sup> | 54 | 4126 |
| HeLa NT cholesterol depleted + TGY<sup>14</sup> | 79 | 2380 |
| HeLa hCD59 KO cholesterol depleted + TGY<sup>14</sup> | 2 | 232 |

As expected, ILY bound well to cholesterol depleted cells, but not to hCD59 KO cells. VLY bound cells in the absence of either cholesterol or hCD59, although more binding was detected on cholesterol depleted cells than hCD59 KO cells. PLY bound well to hCD59 KO cells but showed reduced binding to cholesterol depleted cells (**Fig 4B**). Since PLY is known to require cholesterol for toxin function, the ∼32% of PE-positive cholesterol depleted cells suggest that cholesterol depletion is not completely effective; however, based on lysis data, the cholesterol depletion is sufficient to block PLY-mediated lysis.

For the TGY variants (**Fig 4A**), the flow cytometry results suggest that TGY^3a^ is non-functional due to a defect in host cell binding: only a small percentage of TGY^3a^-exposed cells were PE-positive. The results also indicate that the reduction in binding is due to the glutamic acid in L2 of TGY^3a^, as there was a significant increase in PE fluorescence for TGY^3a_E454G^-incubated cells compared to TGY^3a^-incubated cells, both in the percentage of PE-positive cells and the MFI. Additionally, L2 of TGY^3a^ seems to mediate interaction with the host cell in a cholesterol-dependent manner, as there is minimal binding of TGY^3a_E454G^ to cholesterol depleted cells. This is consistent with previously published findings, which show that L2 inserts into host cell membranes in a cholesterol-dependent manner (32, 33, 35). In hCD59 KO cells, TGY^3a_E454G^ bound ∼60% of the cells, although the significantly lower MFI (compared to TGY^3a_E454G^-incubated control cells) suggests fewer toxin units per cell. This finding suggests that TGY^3a_E454G^ may bind to cholesterol before interacting with hCD59, but that hCD59 is still required for pore formation and lysis, possibly by playing a role in oligomerization – the lower MFI could indicate a lack of concentration of toxin monomers on the surface that occurs during oligomerization.

Of note, TGY^14^ and TGY^3a_E454G^ have similar lysis curves but differentially bind to cholesterol depleted cells: TGY^14^ binds to a significantly higher percentage of cholesterol depleted cells than TGY^3a_E454G^ (**Fig 4A**). This could imply a higher affinity for hCD59 or a higher rate of oligomerization. Furthermore, even though TGY^14^ can bind to a large percentage of cholesterol depleted cells and both functional TGY variants are able to bind to ∼50-60% of hCD59 KO cells, neither of those cell lines are lysed by these toxins (**Fig 3A and 3B**). Taken together, these data suggest that differences in binding ability alone do not account fully for the lysis phenotypes of TGY variants and that subsequent steps of pore formation also play key roles in lysis differences.

### Functional TGY requires both cholesterol and hCD59 for oligomerization

We next investigated the oligomerization of TGY on the host cell surface. HeLa cells were incubated with toxins on ice, before the cells were lysed and prepared for SDS-AGE (20). Glutaraldehyde was used as a cross-linker to preserve oligomer complexes and prevent them from breaking down during sample preparation. For comparison, similar analyses were also carried out with ILY, VLY and PLY.

The resulting western blots demonstrate oligomer “streaks” for ILY and VLY on HeLa NT and HeLa hCD59 KO cells (**Fig S3A**). PLY oligomerization was only evident in both cell lines at higher toxin concentrations (**Fig S3B**), which was unsurprising as the lower PLY concentration causes only ∼10-15% cell death in HeLa NT cells. Oligomerization was not detectable for ILY, VLY and PLY on cholesterol depleted cells (**Fig S3C**).

SDS-AGE analysis with TGY showed visible oligomers for both functional TGY variants (TGY^14^ and TGY^3a_E454G^) on control cells but a lack of visible oligomers for TGY^3a^. Moreover, depleting host cells of either cholesterol or hCD59 abrogated oligomerization for both TGY^14^ and TGY^3a_E454G^ (**Fig 5**). Overall, our data indicate that TGY is dependent on both cholesterol and hCD59 for binding as well as oligomerization.

**Figure 5:**
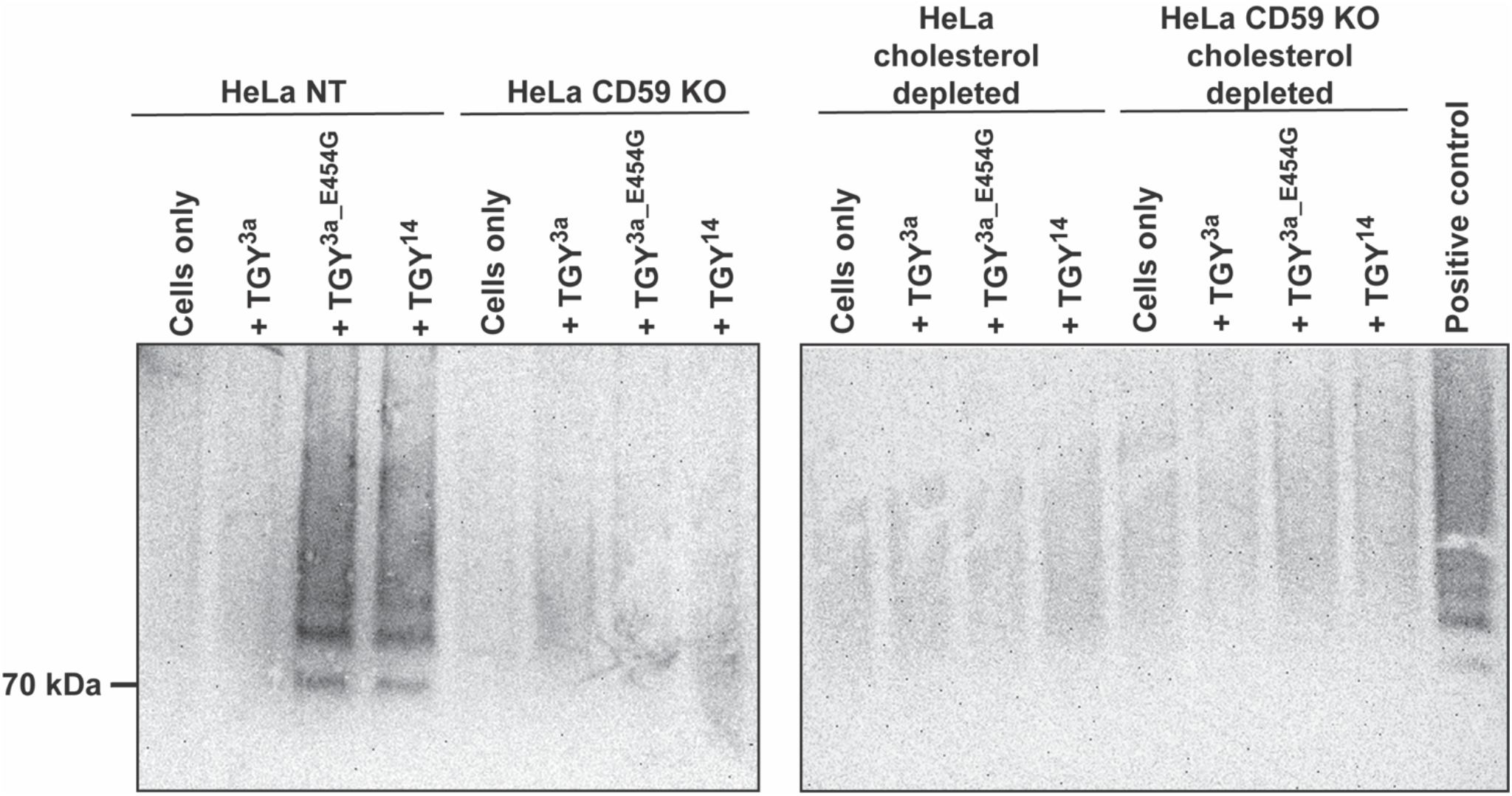
Functional TGY is dependent on both cholesterol and hCD59 for oligomerization on host cell surface. Oligomerization of TGY variants as assessed by SDS-AGE analysis. Cells were incubated with the indicated toxins on ice before being prepared for western blotting. 0.01% glutaraldehyde was used as a cross-linker to preserve oligomers during sample preparation. Western blot membrane was probed with an anti-His-tag HRP-conjugated antibody.

### hCD59-dependent CDCs show increased lysis activity, dependent on host cell hCD59 expression levels

We observed that the concentrations of hCD59-dependent CDCs ILY, VLY and TGY required for lysis differ from those of the non-specific CDC PLY. To investigate how hCD59-dependency affects the overall lytic ability of CDCs, we tested various CDCs for lysis on HeLa cells (**Fig 6A and 6B**). In the non-hCD59-dependent subset, we tested PLY and inerolysin (INY) (38). In the hCD59-dependent subset, we tested ILY, VLY, TGY^3a^, TGY^3a_E454G^ and TGY^14^. To quantify lysis, nonlinear regression analysis was performed on the data to obtain IC_50_ values for each toxin (**Table 2**). All functional hCD59-dependent toxins had lower IC_50_ values than non-hCD59-dependent toxins on HeLa NT control cells. VLY did not behave in a fully hCD59-dependent manner, lysing cells in the absence of hCD59 expression but with lower efficiency (i.e. higher IC_50_ value), as previously indicated in the literature (36). Quantitatively, VLY lyses HeLa control cells at concentrations comparable to ILY (a hCD59-dependent CDC), and HeLa hCD59 KO cells at concentrations similar to PLY and INY (non-hCD59-dependent CDCs) (**Table 2**). These data indicate the possibility that VLY may form pores in a manner similar to either hCD59-dependent CDCs or non-hCD59-dependent CDCs, depending on the characteristics of the target cell. In addition, our data raise the possibility that hCD59-dependency, while restricting a CDC to a single species, might increase lysis efficiency on some target cells.

**Figure 6:**
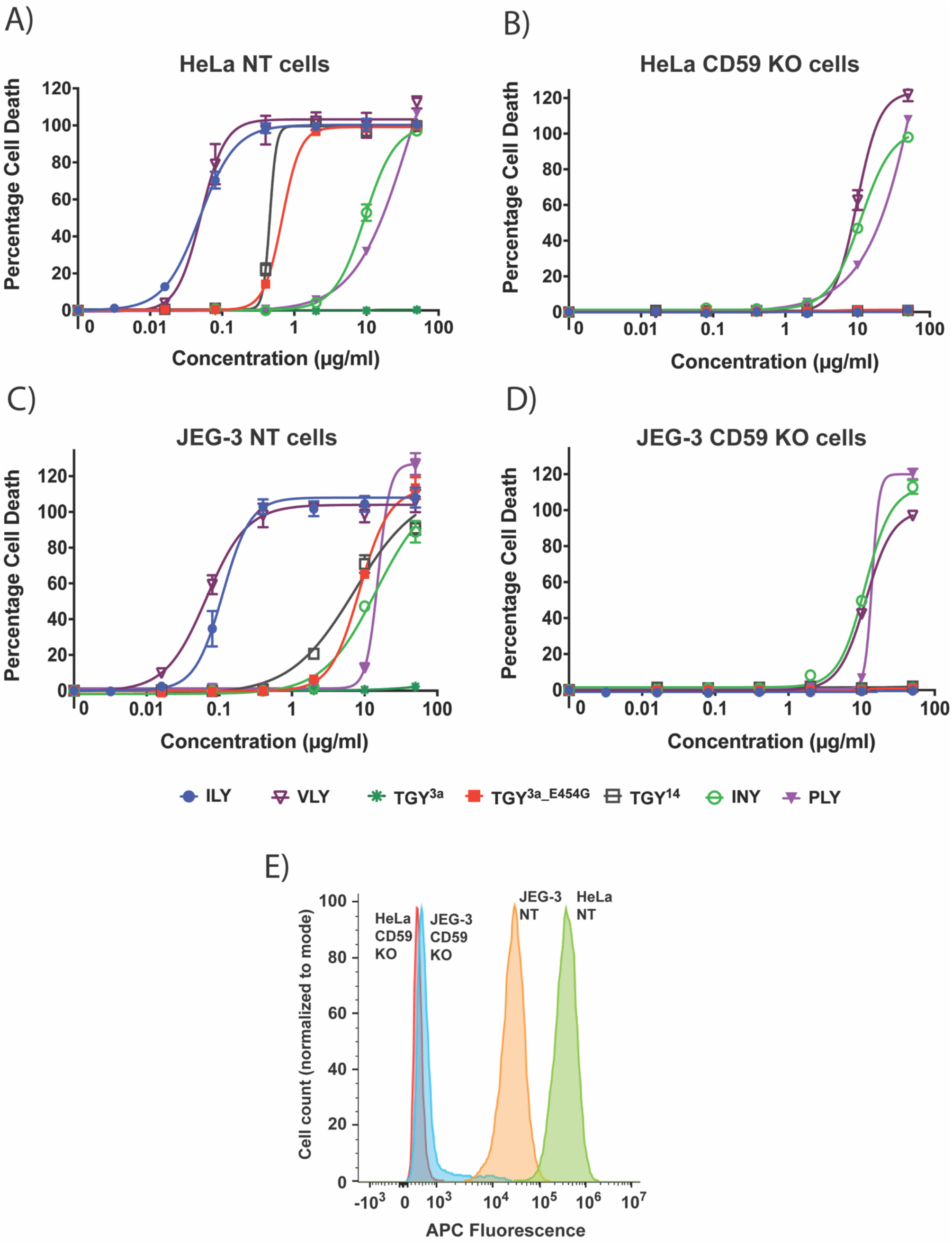
hCD59-dependency coupled with high hCD59 expression increases CDC ability to lyse cells at lower concentrations. (**A-B**) Percentage cell death in HeLa NT control cells and HeLa hCD59 KO cells when exposed to various full-length CDCs, both hCD59-dependent and non-hCD59-dependent. Cells were incubated with toxins for 1.5 hours and cell death measured by an LDH-release cytotoxicity assay. Each point is the mean of 3 replicates, and error bars represent ±SD. (**C-D**) Percentage cell death in JEG-3 NT control cells and JEG-3 hCD59 KO cells when exposed to various full-length CDCs, both hCD59-dependent and non-hCD59-dependent. Cells were incubated with toxins for 1.5 hours and cell death measured by an LDH-release cytotoxicity assay. Each point is the mean of 3 replicates, and error bars represent ±SD. (**E**) Flow cytometry analysis of hCD59 expression on the cell surface for HeLa cells and JEG-3 cells. Cells were incubated with a primary anti-hCD59 mouse IgG2a antibody and a secondary anti-mouse IgG2a AF647 antibody. Histogram x-axis shows fluorescence intensity of the APC-conjugated secondary antibody for each cell population. Histogram y-axis shows cell count normalized to mode. Abbreviations: ILY, intermedilysin; VLY, vaginolysin; TGY, tigurilysin; INY, inerolysin; PLY, pneumolysin.

**Table 2:**
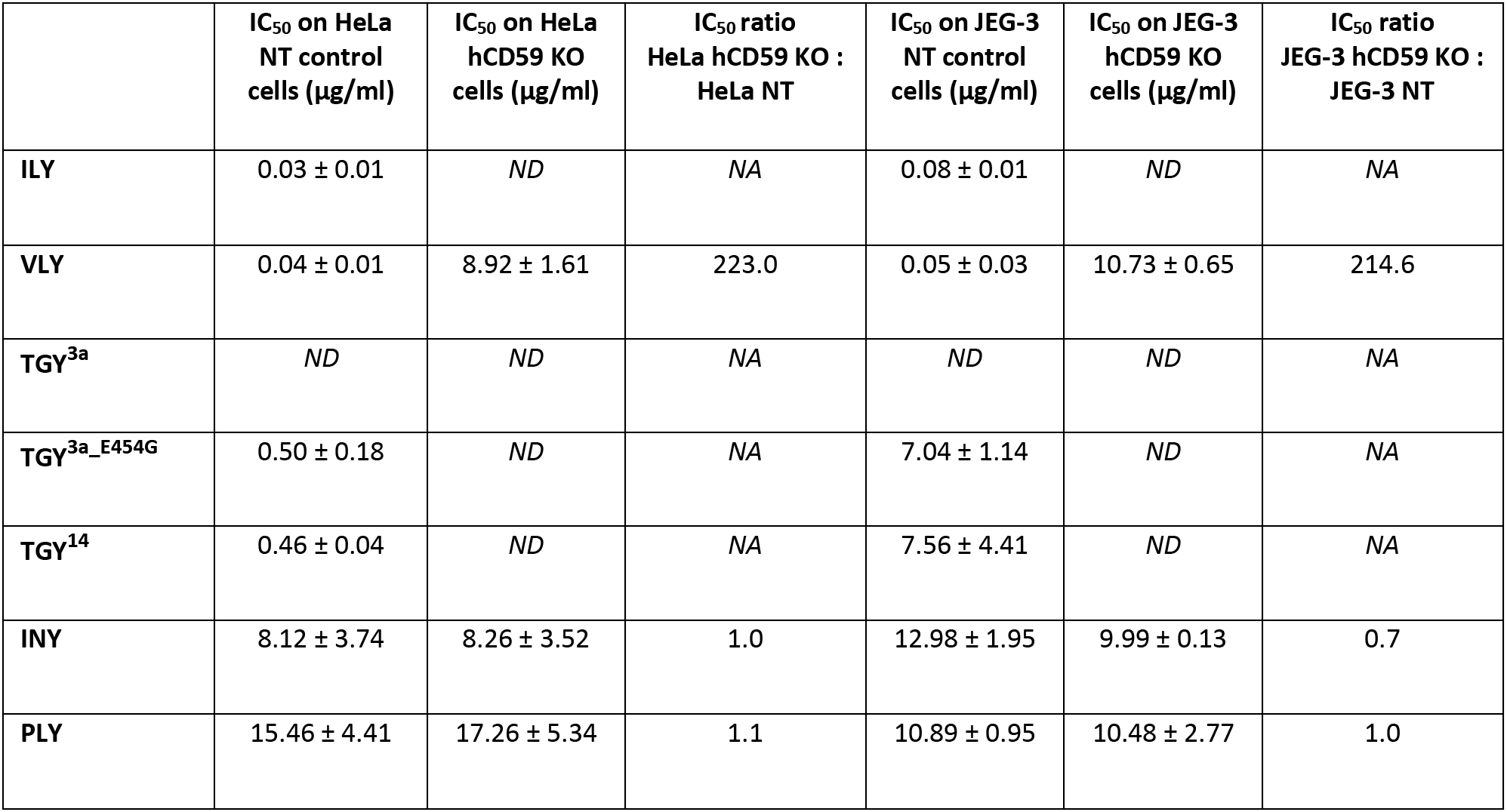
IC_50_ values of CDCs tested for lysis on HeLa and JEG-3 cells. Full-length CDCs were tested for lysis on the indicated cell lines at a range of 0.08 µg/ml to 50 µg/ml, and IC_50_ values indicate the concentrations required for 50% lysis on each cell line. Each IC_50_ value average with standard deviation is calculated from 3 experimental repeats. “ND” indicates where the CDC did not cause any cell lysis; “NA” indicates where the IC_50_ ratio could not be calculated.

To investigate if these lysis results are relevant to other human cell lines, we repeated the LDH release assays with various CDCs on JEG-3 cells (**Fig 6C-6D**). Once again, hCD59-dependent toxins (except for TGY variants) had lower IC_50_ values than non-hCD59-dependent toxins on JEG-3 NT control cells. However, ILY consistently lysed JEG-3 NT cells at IC_50_ values ∼2.5 times higher than HeLa NT cells, and TGY variants lysed JEG-3 NT cells at ∼14-16 times higher IC_50_ values than HeLa NT cells (**Table 2**).

We hypothesized that the difference in lysis efficiency of hCD59-dependent CDCs on two different human cell lines could be affected by the amount of hCD59 expressed on the surface of these cells. To investigate this, we analyzed the cell surface expression of hCD59 on HeLa cells and JEG-3 cells using flow cytometry (**Fig 6E**). The results show that HeLa cells express ∼15 times more hCD59 than JEG-3 cells. These data suggest that changes in hCD59 expression levels between different human cell lines can potentially affect the efficiency of hCD59-dependent CDC pore formation and lysis.

### hCD59-dependency results in increased efficiency of lysis by CDCs

Since Domain 4 is the key player that controls toxin specificity (16–18), we created hybrid CDCs to isolate the role of hCD59-dependency in lysis efficiency. To do this, the Domain 4 of five different CDCs was attached to Domains 1-3 of PLY, a non-hCD59 dependent CDC, resulting in the following: PLY-INY, PLY-ILY, PLY-VLY, PLY-TGY^3a^, PLY-TGY^3a_E454G^ and PLY-TGY^14^.

Lysis caused by each of the hybrid CDCs was measured in HeLa NT and HeLa hCD59 KO cells. The presence of a functional hCD59-dependent Domain 4 increased lysis efficiency of PLY-ILY and PLY-VLY when compared to full-length PLY, while the presence of a non-functional Domain 4 (PLY-TGY^3a^) rendered the hybrid inactive. PLY-TGY^3a_E454G^ and PLY-TGY^14^ also demonstrated a decrease in IC_50_ compared to full-length PLY, albeit less than PLY-ILY and PLY-VLY (**Fig 7A and Table 3**). All hybrid toxins exhibited hCD59-dependency as governed by each Domain 4 (**Fig 7B**). PLY-INY (with a non-hCD59-dependent Domain 4) exhibited a slight decrease in IC_50_ compared to full-length PLY, which could be expected considering full-length INY has a lower IC_50_ than PLY (**Table 3**). PLY-VLY notably retained the ‘intermediate’ nature of its Domain 4 parent: it lysed HeLa NT control cells at low concentrations, and HeLa hCD59 KO cells at concentrations comparable to non-hCD59-dependent PLY, as evidenced by its IC_50_ values. (**Fig 7A**, **7B and Table 3**). Overall, our data suggest that in most cases, while hCD59-dependency restricts CDC targets to a single species, it increases the efficiency of pore formation and lysis for those toxins.

**Figure 7:**
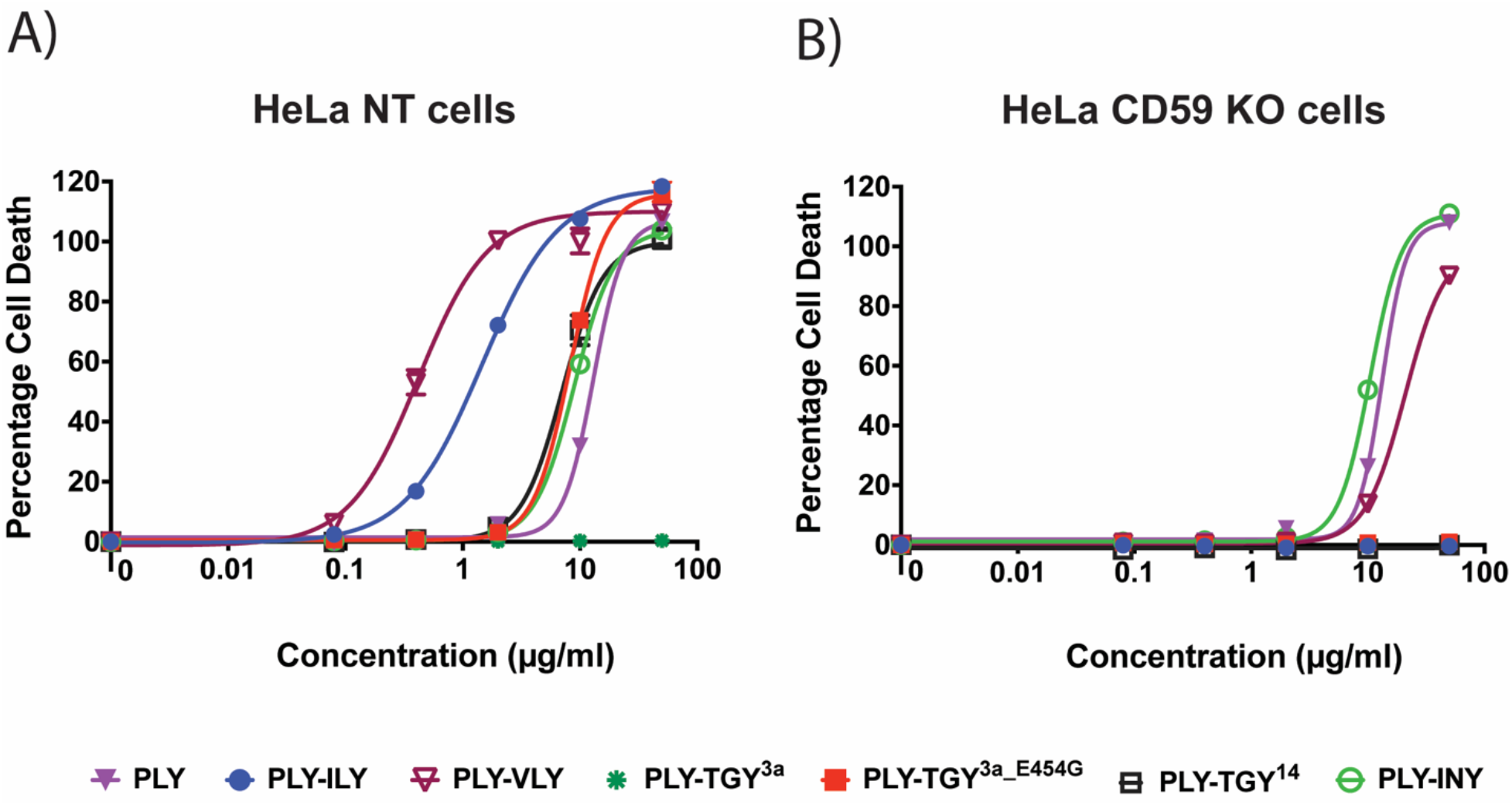
Incorporating a hCD59-dependent Domain 4 increases CDC ability to lyse cells at lower concentrations. (**A-B**) Percentage cell death in HeLa NT control cells and HeLa hCD59 KO cells when exposed to various PLY-hybrid CDCs, with both hCD59-dependent and non-hCD59-dependent D4s. Cells were incubated with toxins for 1.5 hours and cell death measured by an LDH-release cytotoxicity assay. Each point is the mean of 3 replicates, and error bars represent ±SD.

**Table 3:** IC_50_ values of hybrid CDCs tested for lysis on HeLa cells. Hybrid CDCs with PLY D1-3 were tested for lysis on the indicated cell lines at a range of 0.08 µg/ml to 50 µg/ml, and IC_50_ values indicate the concentrations required for 50% lysis on each cell line. Full-length PLY IC_50_ values are included for comparison. Each IC_50_ value average with standard deviation is calculated from 3 experimental repeats. “ND” indicates where the CDC did not cause any cell lysis; “NA” indicates where the IC_50_ ratio could not be calculated.

|  | IC<sub>50</sub> on HeLa NT<br>control cells<br>(µg/ml) | IC<sub>50</sub> on HeLa<br>hCD59 KO cells<br>(µg/ml) | IC<sub>50</sub> ratio<br>HeLa hCD59 KO : HeLa<br>NT |
| --- | --- | --- | --- |
| PLY | 15.46 ± 4.41 | 17.26 ± 5.34 | 1.1 |
| PLY-ILY | 1.82 ± 0.69 | ND | NA |
| PLY-VLY | 0.67 ± 0.26 | 37.60 ± 16.80 | 56.1 |
| PLY-TGY<sup>3a</sup> | ND | ND | NA |
| PLY-TGY<sup>3a_E454G</sup> | 9.48 ± 0.79 | ND | NA |
| PLY-TGY<sup>14</sup> | 11.26 ± 3.73 | ND | NA |
| PLY-INY | 8.66 ± 0.53 | 9.69 ± 0.44 | 1.2 |

## DISCUSSION

Cholesterol dependent cytolysins (CDCs) act as factors in pathogenesis for a wide range of disease-causing bacteria. Understanding their mechanism of action is a key step towards forming therapeutic strategies against such bacteria. Our study adds to the growing knowledge regarding CDCs by characterizing a novel hCD59-dependent CDC (TGY) produced by *Streptococcus oralis subsp. tigurinus*, a bacterial species implicated in various infections (25, 26). Our data demonstrate that of the two TGY variants, TGY^3a^ is non-functional due to a defective L2, which prevents it from binding to host cells and oligomerizing. Furthermore, TGY is fully hCD59-dependent and fully cholesterol dependent, setting it apart from other known CDCs which are generally grouped into categories based on hCD59-dependency (37, 39) but none are known to be equally dependent on both cholesterol and hCD59 for lysis. Overall, TGY seems to exhibit an atypical pore forming mechanism compared to CDCs such as PLY, VLY and ILY.

Our flow cytometry data investigated host cell binding for various CDCs. Binding data with PLY showed that cholesterol depletion for our cells was not complete, with ∼30% cells still being bound by PLY (a fully cholesterol dependent CDC). However, the large shift in PE fluorescence for PLY on control cells versus cholesterol depleted cells, coupled with the PLY lysis data, indicated the cholesterol depletion is sufficient to abolish lysis by nonspecific CDCs. For ILY and VLY, ∼100% of the cholesterol depleted cells are still bound by the toxins but there is a reduction in MFI. This could be due to reduced oligomerization, as fewer oligomers would mean less toxin detectable on the cell surfaces.

TGY^3a^ appears to be non-functional largely due to a defect in binding to host cells. TGY^3a_E454G^ did not bind well to cholesterol depleted cells but could partially bind hCD59 KO cells. This is unlike ILY, which is thought to bind hCD59 before cholesterol (18, 22) and, in our experiments, bound well to cholesterol depleted cells but not to hCD59 KO cells. Notably, TGY^14^ bound to cholesterol depleted cells much better than TGY^3a_E454G^, though the reason for this aberration is unclear.

We conducted SDS-AGE to visualize oligomerization of CDCs on host cell surfaces. Previously, hCD59 dependent CDCs have been shown to require either cholesterol or hCD59 for oligomerization, but not both factors together: ILY can oligomerize on cholesterol depleted cells but requires hCD59 for oligomerization (22, 34, 40), while VLY has been shown to oligomerize on liposomes lacking hCD59 (36). Our oligomerization assays demonstrated no visible oligomers for ILY and VLY in the absence of either hCD59 or cholesterol. The oligomerization data for ILY and VLY correspond to our binding data, which suggested loss of oligomers on cholesterol depleted cells as discussed above. It is relevant to note, however, that all binding and oligomerization data was based on a toxin concentration of 5 μg/ml, which was sufficient for lysis of relevant cell lines when cells are incubated at 37°C for 1.5 hours, but might not be enough for oligomerization when cells are incubated for a shorter time period on ice (as during sample preparation for flow cytometry and SDS-AGE). Therefore, despite ILY and VLY lysing cholesterol depleted cells, and VLY lysing hCD59 KO cells (**Fig 3 and 4**), we observe reduced binding and no oligomerization.

SDS-AGE with TGY variants showed that both hCD59 and cholesterol are important for oligomerization of functional TGY. Consequently, even though TGY^3a_E454G^ bound to about half of hCD59 KO cells, and TGY^14^ bound partially to both cholesterol depleted cells and hCD59 KO cells, no lysis was seen on these cell lines, likely due to a lack of sufficient oligomerization.

Overall, our data demonstrate that TGY residue 454 significantly affects the host cell binding, modulating toxin activity. Furthermore, both cholesterol and hCD59 are required for oligomerization. Our data also imply that the pore forming mechanism of TGY and its interactions with hCD59 and cholesterol potentially vary from those of other hCD59-dependent CDCs. It is prudent to note that the idea of an atypical CDC pore-formation mechanism is not entirely unique. ILY and VLY (both hCD59-dependent CDCs) bind hCD59 differently: VLY has a lower affinity for hCD59 than ILY, and its complex with hCD59 has a different conformation (18). The UDP of VLY can also adopt two different conformations upon interacting with host cell membrane, whereas ILY UDP consistently adopts one orientation (18). The CDC arcanolysin (ALN) displays varying specificity for different mammalian cell lines in a non-hCD59-dependent manner (41). There has also been recent evidence that CDCs might interact with glycans or heparan sulfates on host cell surfaces (42–44). It is therefore possible that host cell factors other than hCD59 and cholesterol, as well as conformational variations in the toxin structure, could play roles in the TGY pore forming mechanism.

The second part of our study aimed to understand the selective pressure behind the evolution of hCD59-dependency in some CDCs. Most CDCs are nonspecific, which means they can target and lyse cells from different species. Conversely, hCD59-dependency restricts CDCs to targeting only human cells. Furthermore, it has been speculated that as hCD59 is present on all human cells, hCD59-dependency would not lead to greater specificity for a particular cell type (45). The data presented in this study demonstrate that hCD59-dependent CDCs like ILY lyse HeLa cells with higher efficiency than non-hCD59-dependent CDCs like PLY and INY. We also showed that CDCs such as VLY behave like intermediate CDCs, lysing cells at low concentrations comparable to ILY when hCD59 is accessible and at higher concentrations comparable to PLY/INY when hCD59 is not available. Additionally, our data showed a difference in hCD59-dependent toxin concentrations needed for lysis of HeLa cells than those needed for JEG-3 cells. This implied that host factors other than the binary presence or absence of hCD59 affected lysis efficiency of these toxins. Flow cytometry data for hCD59 expression on the surface of HeLa and JEG-3 cells revealed a significantly lower amount of hCD59 expressed on JEG-3 cells. The fact that higher concentrations of hCD59-dependent toxins are required to lyse JEG-3 cells leads to the theory that while all human cells express hCD59, differing levels of hCD59 expression could allow hCD59-dependent CDCs to preferentially target some cell types over others in the human body.

Understanding CDC pore-formation is an important step towards developing strategies that can help mitigate PFT-driven bacterial virulence. To this end, our study adds to the growing knowledge around human-specific CDCs by investigating the function of a novel hCD59-dependent CDC. Furthermore, our comparative functional analysis on a range of CDCs endeavors to help us understand the development of human-specificity within this broad family of pore forming toxins.

## MATERIALS AND METHODS

### Cell Culture

HeLa cells (ATCC #CCL-2) and JEG-3 cells (ATCC #HTB-36) were cultivated in EMEM medium supplemented with 10% heat-inactivated fetal bovine serum (FBS) and penicillin/streptomycin. For cells transduced with lentivirus to create NT control cells and hCD59 KO cells, the media was further supplemented with 0.9μg/ml puromycin. Cells were grown in humidified incubators at 37°C and 5% CO_2_.

### Generation of Knockout Cell Lines

NT control and hCD59 KO cell lines were generated as previously described (31). Briefly, single guide RNAs (sgRNAs) were ligated into an empty pLentiCRISPR plasmid backbone (gifted by Dr Feng Zhang; *Addgene plasmid #52961* (46)) and packaged into lentivirus using XtremeGENE 9 DNA Transfection Reagent. Cells were transduced with the lentivirus using spinfection. To generate single cell clones from polyclonal KO cell lines, limiting dilutions were performed.

### Cell Viability Assays (LDH)

LDH Cytotoxicity assay (*Sigma-Aldrich 11644793001*) to measure lysis was conducted according to the manufacturer’s instructions. Briefly, 2.0 x 10^4^ HeLa cells were seeded per well in a 96-well plate overnight. For cholesterol depletion, cells were incubated with 4 mg/ml methyl-ß-cyclodextrin (MßCD) for 1 hour and washed with PBS. Cells were incubated in appropriate dilutions of toxin in fresh medium for 1.5 hours. 100 μl of the supernatant was transferred to a new 96-well plate and 100 μl of Reaction mixture consisting of dye solution and catalyst (prepared according to the manufacturer’s protocol) added. Plates were incubated in the dark for 30 mins. Absorbance of samples was measured at 492nm.

### Recombinant Toxins and Hybrid Toxins

Hybrid toxins PLY-TGY^3a^ and PLY-TGY^14^ were created using Gibson assembly. Primers were designed to isolate PLY Domains 1-3 and the Domain 4 of TGY variants using NEB Builder tool, with overhangs to ensure the alignment of each Domain 4 with PLY Domain 3. The primers were used to PCR amplify the relevant fragments, which were then gel-extracted. A pET28a vector was double-digested using NdeI and XhoI, and gel-extracted. Gibson assembly was carried out with the digested pET28a vector, PLY Domains 1-3, and TGY Domain 4. The reaction product was transformed into NEB 5-alpha cells and minipreps prepared, before being sequenced using T7 primers.

Recombinant PLY, INY, ILY, VLY, PLY-ILY, PLY-VLY, and PLY-INY were prepared as described (38). Full-length and hybrid toxins (TGY^3a^, TGY^14^, TGY^3a_E454G^, TGY^3a_E454A^, TGY^3a_I544L^, PLY-TGY^3a^, PLY-TGY^3a_E454G^ and PLY-TGY^14^) were purified as follows. Briefly, the gene encoding each toxin was codon-optimized for *E. coli*, cloned into pET28a, and transformed into *E. coli* T7Iq. Protein expression was induced by the addition of IPTG. Cells were lysed with a lysis buffer (50 mM NaH_2_PO_4_, 300 mM NaCl, 10 mM imidazole). The toxin was purified from the lysate using HisTrap columns with an FPLC unit into elution buffer (50mM NaH_2_PO_4_, 300mM NaCl, 250mM imidazole), then buffer exchanged into PBS using Amicon Ultra-4 filters. The concentration of each toxin was measured using a Bradford Assay.s

### Site-Directed Mutagenesis

For mutagenesis of TGY residue 454 and 544, the New England Biolabs Q5 Site Directed Mutagenesis kit (*NEB E0552S*) was used according to the manufacturer’s protocols. Briefly, the NEBaseChanger tool was used to generate primers to make the relevant substitutions. These primers were used to PCR amplify the mutated gene incorporated into a pET28a vector, and the PCR product transformed into NEB 5-alpha cells for amplification. Minipreps were prepared and sequenced to confirm the mutations, before transformation into T7Iq cells for protein purification.

### Flow cytometry

Flow cytometry with His-tagged toxins was carried out using direct staining. 1.0 x 10^6^ cells were washed and incubated with 5 μg/ml of each toxin for 10 minutes on ice. Cells were washed with PBS three times and resuspended in 100 μl of ice-cold PBS + 10% FBS + 1% sodium azide. Phycoerythrin (PE) anti-His tag antibody (*BioLegend 362603*) was added to the cell suspension at a dilution of 1:20 and incubated for 30 minutes on ice in the dark. Cells were washed with ice cold PBS, and resuspended in 300 μl of ice-cold PBS + 3% BSA + 1% sodium azide. Flow cytometry to visualize hCD59 expression was carried out using indirect staining as described previously (31). Stained cells were visualized on a CytoFlex Analyzer flow cytometer, and the data analyzed using FlowJo software (version 10.8.1). Cells were gated as follows: live cells using forward scatter area and side scatter area; single cells using forward scatter area and forward scatter height; fluorescence-positive cells using PE-fluorescence intensity or AF647 fluorescence intensity, and forward scatter height. Samples contained at least 10,000 gated live cells.

### Oligomerization western blot

To visualize CDC oligomers, SDS-AGE analysis was carried out as described (20). Briefly, 1.0 x 10^6^ cells were washed then incubated with 5 μg/ml of each toxin (or 30 μg/ml for PLY where indicated) for 10 minutes on ice. Cells were washed twice with PBS and once with HEPES/NaCl buffer (20 mM HEPES at pH 7.5, 150 mM NaCl), before being resuspended in 40 μl of 0.01% glutaraldehyde in HEPES/NaCl as a cross-linker and incubated at 37°C for 15 minutes. 0.1M Tris (pH 8.0) was added to stop the cross-linking reaction. Tris-Glycine SDS Sample buffer (*LC2676; ThermoFisher*) was added to the cell solution at a 1:1 ratio and the mixture incubated at 95°C for 20 minutes to lyse the cells.

The lysate was loaded onto a 1.5% agarose gel. Proteins from the gel were transferred to a polyvinylidene difluoride (PVDF) membrane using semi-wet transfer, blocked in 5% nonfat dry milk in Tris-buffered saline (TBS) plus 1% Tween-20, and incubated with 6x His-tag monoclonal primary antibody (*MA1-135; ThermoFisher*) overnight at a dilution of 1:750. The membrane was washed with TBS plus 1% Tween-20, and incubated with goat anti-mouse IgG secondary antibody (*31430; ThermoFisher*) for 1 hour at a 1:10,000 dilution. The membrane was then washed with several changes of TBS plus 1% Tween-20 for 3 hours before being imaged on iBright CL1000 using Pierce ECL Western Blotting Substrate.

## ACKNOWLEDGEMENTS

This work was supported by grant R01 AI155476 to AJR.

