## Supplementary figures and images for "Characterization of Tigurilysin, a Novel Human CD59-Specific Cholesterol-Dependent Cytolysin, Reveals a Role for Host Specificity in Augmenting Toxin Activity"

### Supplemental Figure 1

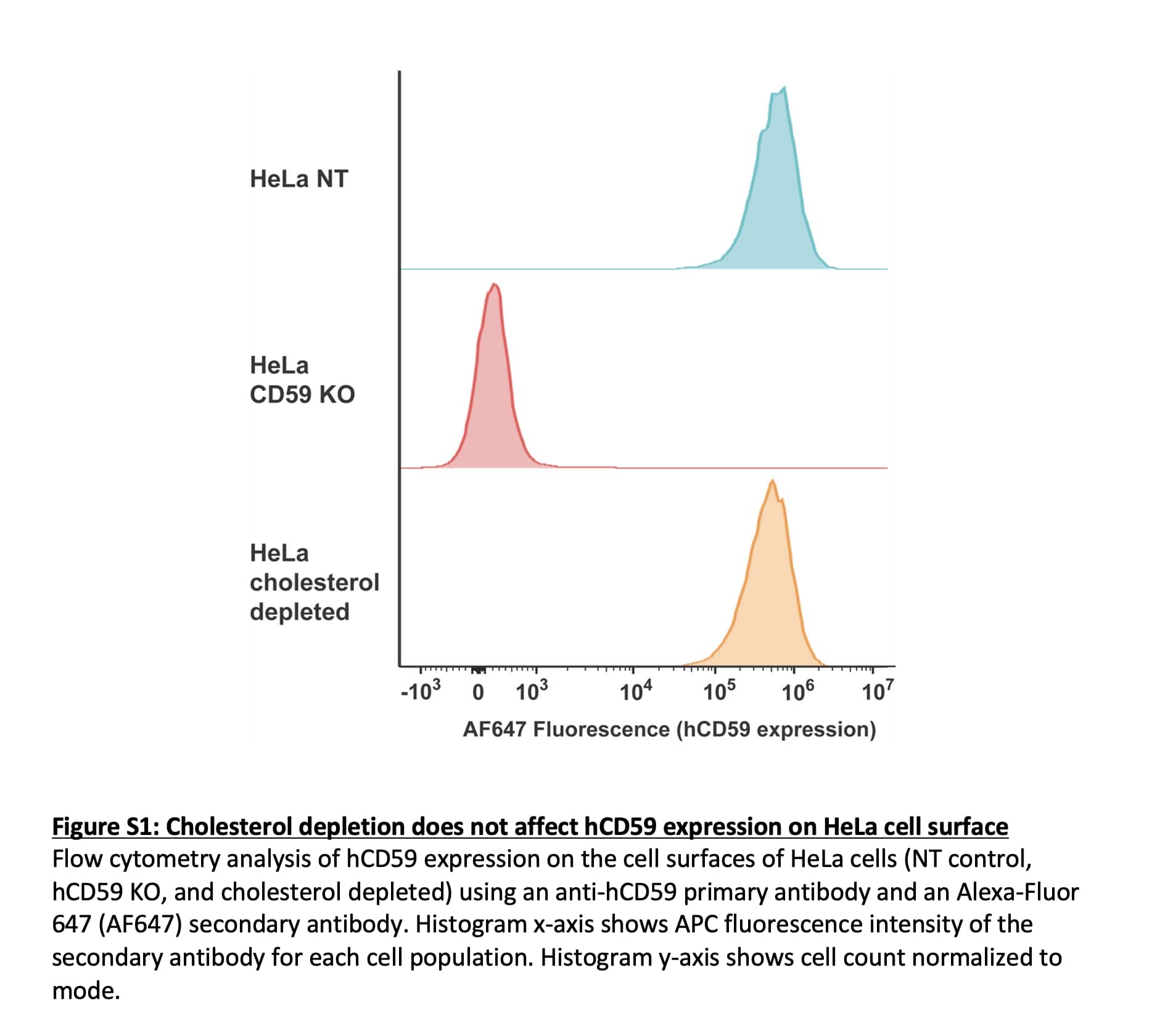

### Supplemental Figure 2

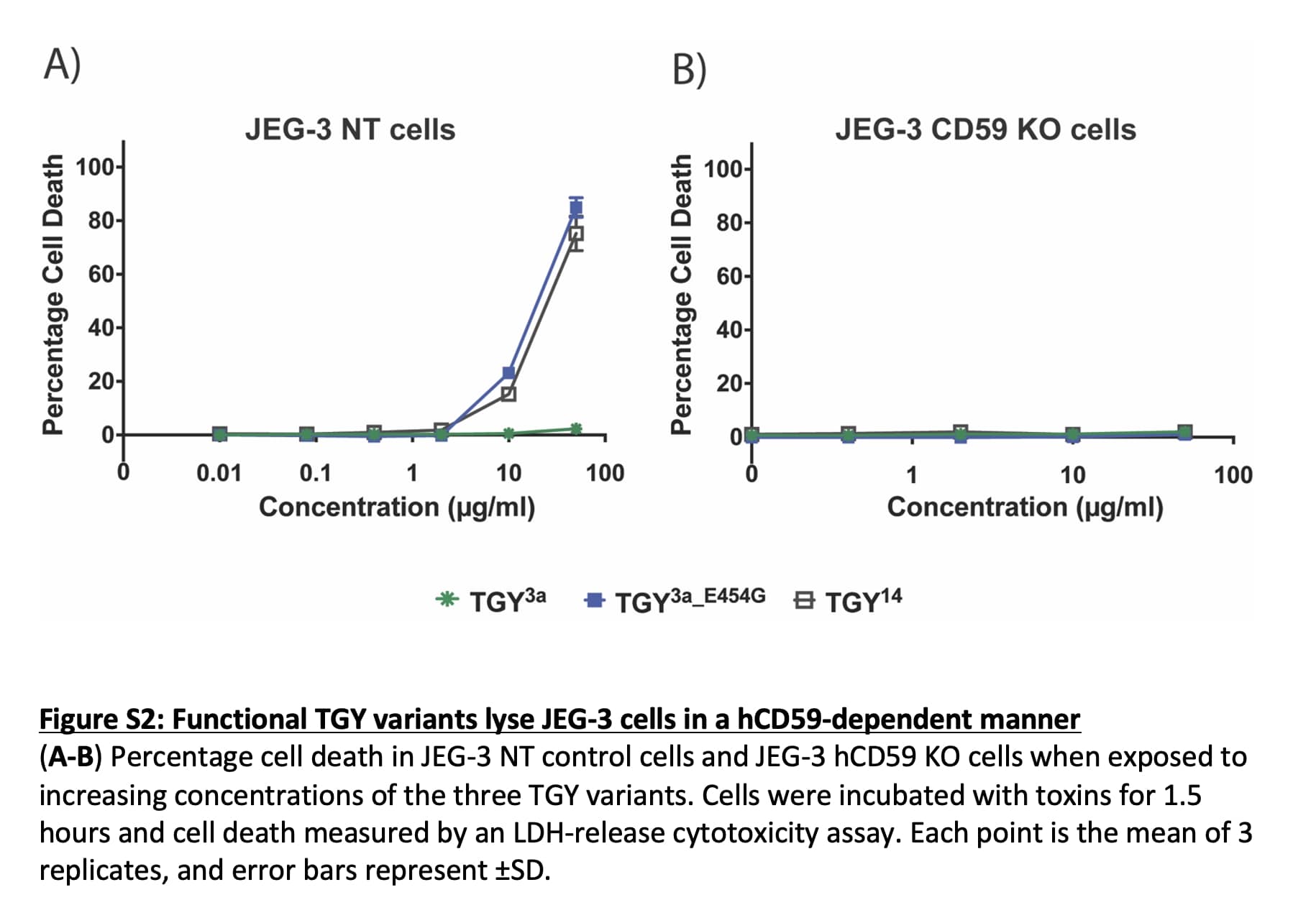

### Supplemental Figure 3

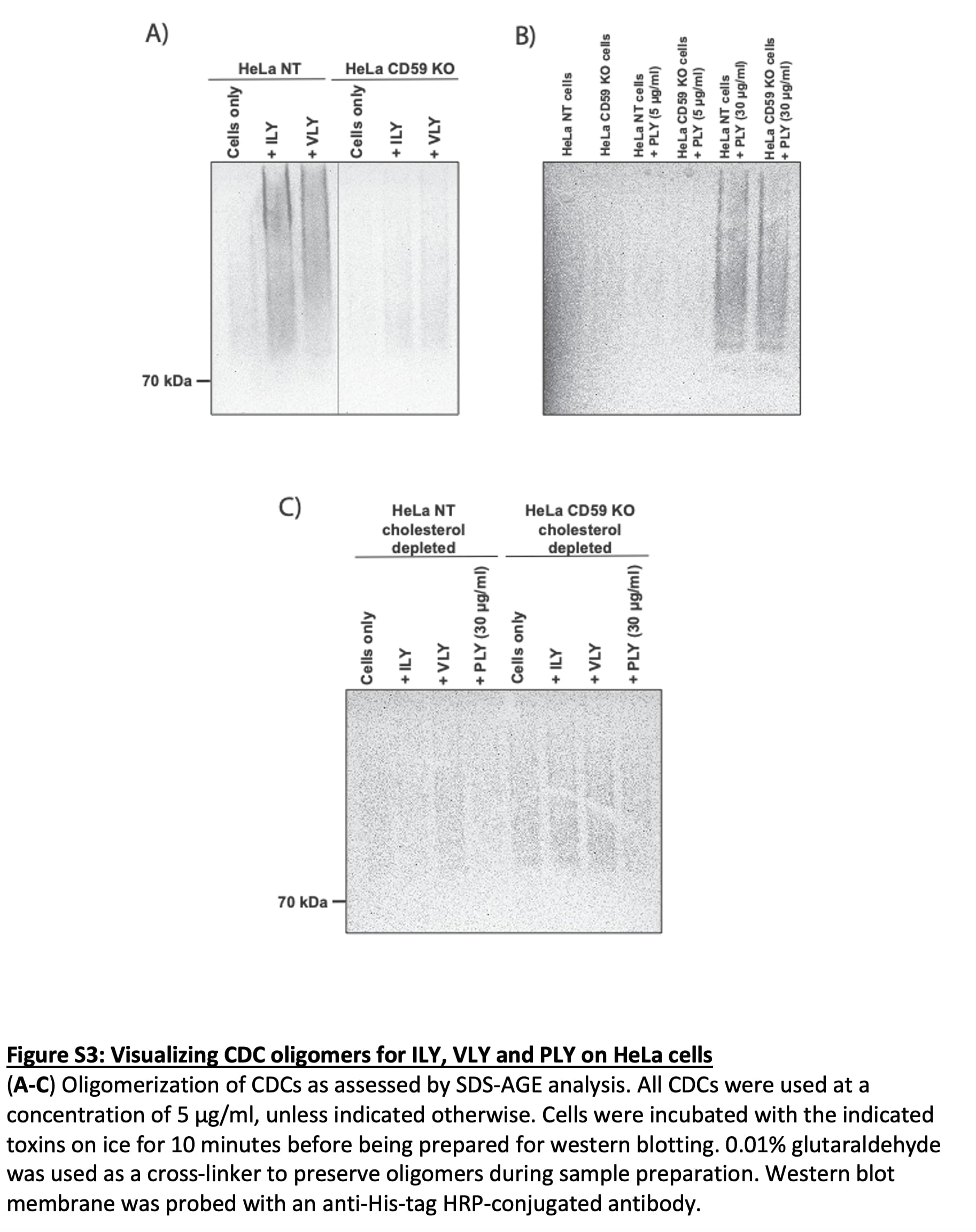
