## Supplemental Table 1 for "Characterization of Tigurilysin, a Novel Human CD59-Specific Cholesterol-Dependent Cytolysin, Reveals a Role for Host Specificity in Augmenting Toxin Activity"

| | Primer name | Sequence (5'-3') |
|---|---|---|
| 1. | PLY_D1-3_F | CTGGTGCCGCGCGGCAGCCATATGGCAAATAAAGCAGTAAATG |
| 2. | PLY_D1-3_R_tgy | CGTTTTTATAAGCTGTAACCTTAGTCTC |
| 3. | TGY_D4_F_ply | GGTTACAGCTTATAAAAACGGCTACCTG |
| 4. | TGY_D4_R | AGTGGTGGTGGTGGTGGTGCTTAGTTGTTTTCAATTTCTTCG |
| 5. | TGY_E454G_F | CACAAGGGTGGTTATGTGGCGC |
| 6. | TGY_E454G_R | ATGCAGATTCAGGTAGCC |
| 7. | TGY_E454A_F | CACAAGGGTGCATATGTGGCG |
| 8. | TGY_E454A_R | ATGCAGATTCAGGTAGCC |
| 9. | TGY_I544L_F | CGGCACCACCCTGCGCCCGAAAT |
| 10. | TGY_I544L_R | TAGTTGGTGATGGTGCGTTTTTG |
| 11. | T7_F | TAATACGACTCACTATAGGG |
| 12. | T7_Term | GCTAGTTATTGCTCAGCGG |

**Table S1: Primers and oligonucleotides**

Primers and oligonucleotides used for construction of hybrids (primers 1-4), site-directed mutagenesis (primers 5-10), and sequencing reactions (primers 11-12).
